# Genetic relatedness cannot explain social preferences in black-and-white ruffed lemurs (*Varecia variegata*)

**DOI:** 10.1101/799825

**Authors:** Andrea L Baden, Timothy H Webster, Brenda J Bradley

## Abstract

Fission-fusion social dynamics are common among a number of vertebrate taxa, and yet the factors shaping these variable associations among subgroup members have not been widely addressed. Associations may occur simply because of shared habitat preferences; however, social ties may also be influenced by genetic relatedness (kinship) or social attraction. Here, we investigate the association patterns of wild black-and-white ruffed lemurs, *Varecia variegata*, in Ranomafana National Park, Madagascar using behavioural, spatial (home range), and genetic data from twenty-four individually identified animals. We collected 40,840 records of group composition over a 17-month period and from this calculated pairwise association indices. We also used ranging coordinates and genetic samples to estimate patterns of spatial overlap and kinship, and then related these measures to patterns of affiliation. From these analyses, we found that dyadic ruffed lemur social associations were generally sparse and weak; that home range overlap was minimal; and that average relatedness within the community was low. We found no evidence that kinship was related to patterns of either spatial overlap or social association; instead, associations were primarily driven by space use. Moreover, social preferences were unrelated to kinship. While home range overlap explained most of the variation seen in social association, some variation remains unaccounted for, suggesting that other social, ecological, and biological factors such as shared resource defense or communal breeding might also play a role in social attraction. Our results further highlight the need to consider individual space use and nuances of species behavior when investigating social preference and social association more generally.

## INTRODUCTION

Animal social systems reflect non-random relationships among neighbouring conspecifics, the content, quality, and patterning of which define their social structure (Hinde, 1976). In fact, there is growing evidence that taxa as diverse as fishes, lizards, birds, cetaceans, and equids exhibit social associations and interactions that are not only non-random, but highly structured (Augusto, Frasier, & Whitehead, 2017; Croft et al., 2012, 2005; Firth et al., 2017; Spiegel, Sih, Leu, & Bull, 2018; Stanley, Mettke-Hofmann, Hager, & Shultz, 2018). Individuals vary in the numbers, strengths, and stabilities of their social ties (Croft et al., 2005; McDonald, 2007; Silk, Alberts, & Altmann, 2006; Silk, Altmann, & Alberts, 2006; Silk et al., 2010a; Stanley et al., 2018), as well as in their preferences for particular social associates (e.g., Cords, 2002; Gero, Gordon, & Whitehead, 2015; Kohn, Meredith, Magdaleno, King, & West, 2015; Mourier, Vercelloni, & Planes, 2012; Perry, 2012; Schülke, Bhagavatula, Vigilant, & Ostner, 2010)—outcomes of which can be evolutionarily significant (Seyfarth & Cheney, 2012; Silk, 2007). For example, the extent, strength, and nature of females’ social relationships have been positively linked to higher fertility (*Macaca mulatta*: Brent et al., 2013; humans: Balbo & Barban, 2014), longer lifespan (*Papio cynocephalus*: Alberts, 2019; Archie, Tung, Clark, Altmann, & Alberts, 2014; *Papio ursinus:* Silk et al., 2010b), and greater offspring survival (*Equus caballus*: Cameron, Setsaas, & Linklater, 2009; *Papio cynocephalus*: Silk, Alberts, & Altmann, 2003; *Papio ursinus:* Cheney, Silk, & Seyfarth, 2016; Silk et al., 2009; *Physeter microcephalus*: Whitehead, 1996). Similarly, males that form strong social bonds have been shown to be more successful at forming coalitions, achieving high rank, and siring more offspring than males with weaker social bonds (e.g., *Chiroxiphia linearis*: McDonald, 2007; *Macaca assamensis*: Schülke et al., 2010; *Pan troglodytes*: Gilby et al., 2013). Nevertheless, while the long-term health and fitness outcomes of sociality have been the focus of much investigation, until recently the factors shaping social preferences have received comparatively less attention.

Several hypotheses have been proposed to explain social association, the most popular being kinship, whereby group members preferentially associate with and, in the case of kin selection, are more likely to direct costly altruistic behaviours toward close genetic relatives (Hamilton, 1964). For instance, many female mammals associate with and exhibit social preferences toward maternal kin (Smith, 2014), a pattern observed in primates in particular (reviewed in Langergraber, 2012), but also in spotted hyaenas (Wahaj et al., 2004), African elephants (Archie, Moss, & Alberts, 2006), dolphins (Frère et al., 2010), and mountain goats (Godde, Côté, & Réale, 2015), among others. Social preferences toward kin can be driven by the benefits of associating with relatives, such as allomaternal care (reviewed in Briga, Pen, & Wright, 2012; Pope, 2000; and in some cases even brood parasitism, Andersson, 2017), reduced aggression and infanticide risk (reviewed in Brown & Brown, 1996; Agrell, Wolff, & Ylonen, 1998), foraging advantages (Griffiths & Armstrong, 2002; Nystrand, 2007), and shared social and ecological knowledge (McComb, Moss, Durant, Baker, & Sayialel, 2001; Salpeteur et al., 2015). However, social associations have also been linked to direct fitness consequences among nonrelatives (Baden, Wright, Louis, & Bradley, 2013; Cameron et al., 2009; G. G. Carter & Wilkinson, 2015; McFarland et al. 2017; Riehl, 2011), suggesting that social preferences can evolve based on direct benefits alone. Moreover, evidence for kin-biased relationships in other species, particularly those with higher fission-fusion dynamics, is weak (e.g., Best, Dwyer, Seddon, & Goldizen, 2014; K. D. Carter, Seddon, Frère, Carter, & Goldizen, 2013; Hirsch, Prange, Hauver, & Gehrt, 2013; Langergraber, Mitani, & Vigilant, 2007, 2009; Moscovice et al., 2017; Wilkinson, Carter, Bohn, & Adams, 2016), raising questions as to the ubiquity of kinship in the formation and maintenance of social bonds.

One complicating factor in determining whether and how kinship shapes social preference is that it can be difficult to decouple social associations due to kinship from associations due to other social and spatial contexts (reviewed in Doreian & Conti, 2012; Wey, Blumstein, Shen, & Jordán, 2008). For example, in some species it is difficult to differentiate whether strong associations are due to strong genetic ties or to other factors, such as sex or age (Lusseau & Newman, 2004; Mourier et al., 2012), habitat utilization (Wiszniewski, Allen, & Möller, 2009), or foraging specializations (Daura-Jorge, Cantor, Ingram, Lusseau, & Simões-Lopes, 2012; Gilby & Wrangham, 2008; Griffiths & Armstrong, 2002; Mitani, Merriwether, & Zhang, 2000). Indeed, patterns of social association have been found to correlate with spatial overlap (i.e., overlapping home ranges) more strongly than, or to the exclusion of, kinship in a number of species (Best et al., 2014; K. D. Carter et al., 2013; Frère et al., 2010; Strickland, Gardiner, Schultz, & Frère, 2014). Moreover, although individuals living in closer physical proximity are more prone to interact (e.g., Clutton-Brock, 1989; Kossinets & Watts, 2006), conspecifics with high levels of spatial overlap may also mediate their interactions by temporally modifying range use, such that animals with highly overlapping ranges may interact minimally, if at all (e.g., temporal avoidance) (Leu, Bashford, Kappeler, & Bull, 2010; Ramos-Fernández, Boyer, Aureli, & Vick, 2009; Strickland et al., 2017). While still uncommon, studies that consider the simultaneous effects of spatial overlap and kinship on association patterns are thus important to understanding the role of social preference in the evolution of social systems and sociality (e.g., Best et al., 2014; K. D. Carter et al., 2013; Frère et al., 2010; Lusseau et al., 2006; Maher, 2009; Podgórski, Lusseau, Scandura, Sönnichsen, & Jȩdrzejewska, 2014; Strickland et al., 2014; Piza-Roca et al. 2019).

Ruffed lemurs (Genus *Varecia*) are moderately-sized frugivores (Baden, Brenneman, & Louis Jr., 2008; Balko & Underwood, 2005) that live in groups (hereafter “communities”) comprising as many as 30 adult and subadult individuals and their offspring (Baden, Webster, & Kamilar, 2016 and references therein). From earlier work, we know that communities are characterized by high fission-fusion dynamics (*sensu* Kummer 1971), with members associating in fluid subgroups (or ‘”parties”) that vary in size, cohesion, membership, and duration (Baden, Webster, & Kamilar, 2016). Within communities, there is striking variation in individual ranging behavior, degree of home range overlap with other community members, and the number and strength of social associations (Baden, 2011; Baden & Gerber, n.d.; Baden et al., 2016; Vasey, 2006). Individuals never use the entire communal territory, instead concentrating their ranging to proportionately smaller areas that overlap with as few as four (25%) to as many as twelve (75%) other community members (Baden & Gerber, n.d.). Association patterns are equally variable, with individuals associating in subgroups with anywhere from one to ten individuals throughout the year (Baden et al. 2016). Overall, associations are generally weak, and animals spend nearly half of their time alone (Baden et al. 2016). Nevertheless, certain individuals appear to preferentially associate and are regularly found together in “core groups” whose members are characterized by both high association strength and home range overlap (Vasey 2006; Morland 1991a,b; Baden 2011). Moreover, studies of microsatellite STR markers and mitochondrial DNA have shown that, while average pairwise relatedness within communities is low (i.e., both sexes disperse: Baden et al., 2014), communities nevertheless contain relatives (e.g, Baden, 2011; Baden et al., 2013). Taken together, these lines of evidence suggest that during subgroup formation community members may be actively choosing social associates, and that these associations may be shaped by a number of ecological, social, and biological factors. Here, we hypothesize (H1) that social associations among members of a black-and-white ruffed lemur (*Varecia variegata*) community are driven by space use and kinship. Specifically, we quantify dyadic measures of association strength, spatial overlap, and relatedness to test the predictions that members with higher home range overlap (P1.1) and relatedness (P1.2) will associate more often than dyads which exhibit disparate range use or are unrelated. Moreover, because previous studies have found a relationship between social preference and kinship, we further hypothesize (H2) that relatedness will drive association preferences (P2.1), with preferred associates being more related than non-preferred dyads.

## METHODS

### Study site and subjects

We collected data from one *V. variegata* community at Mangevo [21°220 6000 S, 47°280 000 E], a mid-elevation (660-1,200 m) primary rainforest site located in the southeastern parcel of Ranomafana National Park, Madagascar (Wright et al., 2012) during 16 months of study (August–December 2007; February– December 2008). At the time of the study, the community included 24 adults and subadults (8 adult females, 11 adult males, 5 subadult males). All individuals in this study were habituated and individually identified via radio-collars or unique collar-tag combinations prior to behavioural observations (see Baden et al., 2016 for specific details; see also Ethical note below). Animals were collared under veterinary supervision following a strict protocol (Glander, 1993). Nineteen infants were born in the 2008 birth season and were present from October to December 2008, when the study ended. Sampling efforts resulted in a total of 4,044 focal observation hours.

### Ethical note

All animal protocols adhere to the guidelines for the treatment of animals for teaching and research recommended by ASAB/ABS (2019). Subjects were habituated to researcher presence as part of a long-term study established in 2005. Since 2005, subjects have been periodically immobilized in the field (approximately once every 2 to 3 years) for application or replacement of radio-transmitter collars, which allow researchers to accurately identify individuals and locate and track them in the field during behavioral observations. Immobilization and sampling procedures adhered to the protocols established by Glander (1993) and have been implemented by Dr. Randall Junge (MS, DVM, Dipl American College of Zoological Medicine, Dipl American College of Animal Welfare), Director of the Prosimian Biomedical Survey Project (PBSP), since 2000. During captures, each lemur is immobilized by remote injection with tiletamine/zolazepam. Animals are examined immediately upon capture, and once determined to be stable they are transported back to the basecamp for examination (not more than 15 minutes). Animals are evaluated again for stability, and monitored by a licensed veterinarian (Dr. Junge) throughout the anesthesia. Biological samples, including blood (no more than 1 ml per kilogram) for genetic analysis and biomedical health assessments, are collected for later analysis. Lemurs are given a balanced electrolyte solution (30 ml) subcutaneously and are monitored until recovery. Once fully recovered, lemurs are released at the site of capture. Anesthetic episodes typically last 1 hour or less. Lemurs that are adequately awake by midafternoon (3 PM) are released the same day. Those that are not adequately recovered are held overnight in cloth bags and released before 10 AM the next morning. As ruffed lemurs are diurnal, releasing animals with residual sedation in the late afternoon could predispose them to predation. This capture, monitoring, sampling and release protocol has been used successfully by PBSP veterinarians for 15 years, on 44 trips to 18 sites throughout Madagascar. Biomedical health assessments by Dr. Junge and population genetic assessments by Baden et al. (2014, 2019) are integral parts of population management, as they are used to identify heath concerns such as introduction of human or livestock parasites to the site (as documented in other parts of Ranomafana National Park) and inbreeding depression. They are also useful for identifying health parameters in a population inhabiting a pristine forest, allowing that information to be used as a benchmark to evaluate this and other populations should they experience epidemic disease or population decline. Observers never interact with subjects outside of the capture period. Utmost care was taken to minimize the impact of our presence on our subjects during captures and subsequent behavioral observations. Observations were conducted noninvasively at a minimum observer distance of 10 m from the focal subject. When in the presence of subjects, observers speak quietly and make efforts to not disturb the individual. No more than four observers are allowed to participate in observations at any one time. Subjects were target for focal observations, at most, twice per month. Permission to conduct research was granted by Madagascar’s National Parks (ANGAP/MNP, #084/07-041/08). Research protocols were approved by and in compliance with Stony Brook University IACUC #2005-20081449.

### Data collection

#### Behavioural monitoring

We collected data during dawn to dusk follows of focal individuals. We located focal subjects at the beginning of each observation period via radio-telemetry and selected new subjects daily. We never sampled focal subjects on consecutive days and every effort was made to follow all subjects at least once per month. If an individual with a collar-tag was located in association with a radio-collared focal individual prior to 10:00 h, this individual became the new focal subject for that observation period. Observational periods ranged in duration between 8 to 11 hours depending on seasonal differences in day length and time needed to locate animals at dawn.

Upon initial contact with the focal individual, we recorded the number and identities of all other individuals present within the subgroup. To do so, one observer remained with the focal individual while the remaining three team members spread out to locate and identify all other members of the subgroup. *A priori*, individuals were considered members of a subgroup only if they were within 50 m of the group center and were seen associating, traveling with and maintaining proximity to members of the subgroup being followed. However, an earlier study found that average group spread in this population ranged between 6 to 15 m, and rarely exceeded 30 m (Baden et al. 2016). Thus, while subgroup membership was operationalized to include all members within 50 m, effective subgroup spread was typically much smaller (see Baden et al., 2016 for details). After initial contact, we monitored subsequent changes in subgroup size, composition (age/sex class, individual identity), and cohesion (i.e., the greatest distance between any two subgroup members), as well as activity state of the focal subject using instantaneous scan sampling techniques collected at 5 min intervals (Altmann, 1974). Sampling efforts resulted in a total of 4,044 focal observation hours, during which time we recorded 40,840 group scans.

We collected simultaneous GPS coordinates at 10 min intervals from as close to the focal individual as possible to document daily individual range use. We recorded spatial coordinates only if estimated positional error was less than 10 m.

#### Genetic sampling

In addition, we collected genetic samples from 38 black-and-white ruffed lemurs from the Mangevo population, including all subjects in this study, during five capture seasons spanning four consecutive years (2005-2008; see Baden, 2011 for details). Sample collection occurred under veterinary supervision and followed a strict protocol outlined by Glander (1993). All capture procedures occurred during non-reproductive seasons in the absence of infants and dependent offspring.

For each individual captured, one of us (ALB) collected approximately 1 ml/kg of whole blood (∼4 cc) from the femoral vein and four 2mm tissue biopsies from ear pinnae. All samples were stored in 5 ml of lysis buffer solution (0.1 M Tris-HCl pH, 8.0, 0.1 M EDTA, 0.01 M NaCl, and 0.5% w/v SDS) at ambient temperature in the field (7 to 21 days) (Longmire, Gee, Hardekopf, Graham, & Mark, 1992). We then banked samples in a −80°C freezer at the Madagascar Biodiversity Partnership headquarters in Antananarivo, Madagascar and subsequently at the Yale Molecular Anthropology Lab in New Haven, CT until later genetic analysis (see below).

### Data analysis

#### Social association

In this study, we define social association as individuals being present in the same subgroup at the same time. We considered social preferences to occur (that is, the dyad was composed of “preferred associates”) when dyads had a significantly stronger strength of association than other dyads in the community (see “Relatedness among preferred associates” below). This species is typically described as being “female-bonded” (Morland 1991a), in line with the definition that to be ‘socially bonded’ requires that multiple, independent types of associations or interactions are significantly higher than expected (Whitehead, 2008). We do not examine social bonds in this study because we only focus on a single measure of association. Finally, we also use the term “social tie” in the network sense (e.g., Wasserman and Faust, 1994; Croft et al., 2008): individuals that have a social tie have a nonzero association index and a “stronger tie” has a higher association index.

Sampling was biased toward radio-collared females; we therefore subsampled our data prior to association analyses. Details are presented in Baden et al. (2016), but briefly, we divided our dataset into monthly periods and according to the sex of the focal individual. Using only scans for which all individuals were known, we randomized point scans and selected X scans to include in the dataset for each sex, where X is 90% of the point scans for the sex with the fewest scans in a given month. This procedure resulted in the inclusion of a total of 11,784 point scans, with equal numbers of scans targeting male and female focal subjects in each month (Baden et al., 2016).

We used SOCPROG 2.9 (Whitehead, 2009) to calculate association indices (AIs) between all pairs (i.e., dyads) of individuals using the “simple ratio” index, which quantifies the proportion of time that two individuals are observed together relative to their total observation time overall. This index is appropriate when individuals are equally likely to be correctly identified (Cairns & Schwager, 1987; Ginsberg & Young, 1992; Whitehead, 2008), which is the case for this population. Given that subgroup composition in this community changes approximately every 90 minutes (Baden et al., 2016), we used a 6-hour sampling interval to reduce autocorrelation among periods. We also removed any subject observed in fewer than 10 sampling periods. Our final dataset (overall dataset) included 8 adult females and 9 adult males totaling 136 dyads observed in over 11,171 point scans.

To examine seasonal variation in associations, we organized the subsetted data from 2008 into three seasons according to female reproductive state as defined by Baden et al. (2013). Using the protocols described above, our final datasets included: 8 females and 9 males over 5,051 scans during the nonbreeding season (January through June); 7 females and 8 males over 3,267 scans during the mating/gestation season (July through September); and 8 females and 6 males over 2,664 scans during the birth/lactation season (October through December). We compared distributions of AIs across seasons using a Kruskal-Wallis test implemented in R (Kruskall and Wallis, 1952).

#### Spatial overlap

Estimates of home range overlap used in this study come from Baden & Gerber (n.d.). Briefly, we calculated home range overlap between all pairs of individuals using a utilization distribution overlap index (UDOI; Fieberg & Kochanny, 2005) implemented in the R package adehabitat (Calenge, 2011). The UDOI is an index of space-use sharing between two utilization distributions (UDs). UDOI values can range from 0 to 1, with a UDOI of 0 indicating no home range overlap and a UDOI of 1 indicating that home ranges are uniformly distributed and have 100% overlap. Values can also be >1 if both UDs are nonuniformly distributed and also have a high degree of overlap. Values <1 indicate less overlap relative to uniform space, whereas values >1 indicate higher than normal overlap relative to uniform space. We calculated four UDOIs for all pairs of individuals: one annual UDOI, and three seasonal UDOIs according to female reproductive state, as for association analyses described above.

#### Relatedness

We genotyped individuals at a suite of 15 polymorphic microsatellite loci (see Baden, 2011 and Baden et al., 2014 for details). We extracted total genomic DNA from blood and/or tissue samples using standard nucleic acid extraction kits (QIAamp DNA Mini Kit; Qiagen) automated on a QiaCube (Qiagen). Extraction procedures followed the manufacturer’s protocols, with the following modification to the tissue extraction procedures: samples were allowed to lyse initially in ASL buffer for 24-48 hours rather than 10 minutes.

We carried out PCR amplifications in a total reaction volume of 25 μl consisting of 2 μl template, 12.5 μl Qiagen HotStar Taq Master Mix, and 10 μM of each primer. Amplification conditions were as follows: initial denaturation at 95 °C for 15 min; 35 cycles of 30 s at 94 °C, 40 s at 54 to 60 °C (see Louis et al., 2005), 1 min at 72 °C, and a final extension of 7 min at 72 °C. The 5′ end of the forward primer was fluorescently labeled, and amplification products were separated and visualized using capillary electrophoresis (ABI 3730xl Genetic Analyzer). We assessed allele sizes relative to an internal size standard (ROX-500) using Gene Mapper software (Applied Biosystems), and scored final genotypes based on multiple independent reactions (Taberlet, 1996). Panels yielded PIsib (Queller & Goodnight, 1989) values of 2.7×10^−5^, demonstrating the very low probability that two individuals would share the same multilocus genotype by chance. We further tested the robusticity of this suite of loci for estimating relatedness with a rarefaction analysis as in Altmann & Alberts (1996) and de Ruiter & Geffen (1998) using the program RE-RAT (http://people.musc.edu/~schwaclh/). We estimated pairwise relatedness among individuals (*r*) following Queller & Goodnight (1989) using the program GenAlEx 6.5 (Peakall & Smouse, 2012). Relatedness was based on allele frequencies derived from a larger population of 38 adult multilocus genotypes (Baden, 2011). Fine scale dyadic relatedness assessments (e.g., distinguishing between full and half-sibs) are not possible in most microsatellite studies, and in fact, the ability to differentiate relatedness disjunctions on such a scale would probably require 30 to 60 microsatellite loci (Stone & Björklund, 2001). We therefore consider “related dyads” to be those with r-values ≥ 0.25; we made no attempt to further distinguish categories or degrees of relatedness.

#### Relationships among social associations, spatial overlap, and relatedness

We used a series of Mantel tests followed by a multiple regression quadratic assignment procedure (MRQAP) to test our first hypothesis (H1), that association strength was driven by spatial overlap and relatedness. We first used Mantel tests to examine whether association indices were independently related to either spatial overlap (P1.1) or relatedness (P1.2), as well as to test for a correlation between spatial overlap and relatedness (Mantel, 1967). We further used a multiple regression quadratic assignment procedure (MRQAP; Dekker, Krackhardt, & Snijders, 2007) to determine relationships between the response variable—association index—and predictor variables—sex, spatial overlap (P1.1), and relatedness (P1.2). For sex, we used a matrix of sex similarity, which used values of 1 (same sex) and 0 (different sex). MRQAP tests each pairwise combination of response and predictor matrices while holding the remaining predictor matrices constant. We performed both Mantel tests and MRQAP for the overall dataset as well as in each of the three seasons using SOCPROG 2.9 (Whitehead, 2009).

#### Relatedness among preferred associates

Studies of other taxa have found that relatedness can shape social preferences, even if it does not structure association among all dyads (Carter et al., 2013; Best et al., 2014). Therefore, in addition to the analyses described above, we used generalized linear models to test our second hypothesis (H2) that relatedness between two individuals in a dyad predicted whether they were “preferred” associates. Here, we define “preferred associates” (we also use “social preference”) as dyads exhibiting significantly higher association indices than other dyads in the community.

We identified preferred associates in our overall dataset (described in “Social association” above) using SOCPROG 2.9’s (Whitehead, 2009) permutation tests based on Bejder et al. (1998) with modifications by Whitehead (2008). We used the “permute associations within samples” option, along with 1000 flips per permutation and 10,000 permutations. We classified dyads as preferred associates if the dyad’s AI was significantly high (one-tailed *α* = 0.05).

We next used logistic regression analysis, implemented using Scikit-learn in Python (Pedregosa et al., 2011) to test if relatedness of a dyad (independent variable; measured using continuous relatedness values calculated as described in “Relatedness” above) predicted whether members in that dyad were preferred associates (P2.1; dependent variable; binary response of “preferred” or “not” based on SOCPROG permutation analyses). To assess significance, we compared the coefficient of the regression model to a “null” distribution of coefficients generated from 10,000 randomizations of the model. During each replicate, we shuffled relatedness values among dyads before building the randomized regression model. Our p-value was calculated as: 1 – (*percentile of observed coefficient in null distribution*). The script in which we implemented this procedure is available at https://github.com/thw17/Varecia_social_preferences.

## RESULTS

### Social association

The average overall association index (AI) was 0.05 ± 0.03 (Baden et al., 2016). While we observed similar mean AIs in each of the three reproductive seasons (nonbreeding = 0.04 ± 0.02; mating/gestation = 0.06 ± 0.03; birth/lactation = 0.07 ± 0.03), the distributions of AIs significantly differed across seasons (Kruskall-Wallis chi-squared = 8.423, df = 2, p = 0.015). Within seasons, individuals also varied substantially in their AIs. Distributions were right skewed, with many values at or near 0 and maximum values as high as 0.82. Average individual AIs (i.e., the average of all of an individual’s AIs) varied from 0.01 to 0.11 (nonbreeding = 0.01-0.08; mating/gestation = 0.01-0.11; birth/lactation = 0.01-0.10), while maximum individual AIs ranged from 0.07 to 0.82 (nonbreeding = 0.08-0.82; mating/gestation = 0.07-0.79; birth/lactation = 0.08-0.82).

### Spatial overlap

Average home range overlap (UDOI) was 0.211 ± 0.357 (Baden & Gerber, n.d.). Some dyads did not overlap at all (UDOI = 0), while the maximum UDOI observed was 1.895 (Figure 1). As with AIs, we observed similar mean UDOIs in each of the three reproductive seasons (nonbreeding = 0.17 ± 0.27; mating/gestation = 0.15 ± 0.29; birth/lactation = 0.20 ± 0.43; Table 1; see also Baden & Gerber n.d.)

**Figure 1.**
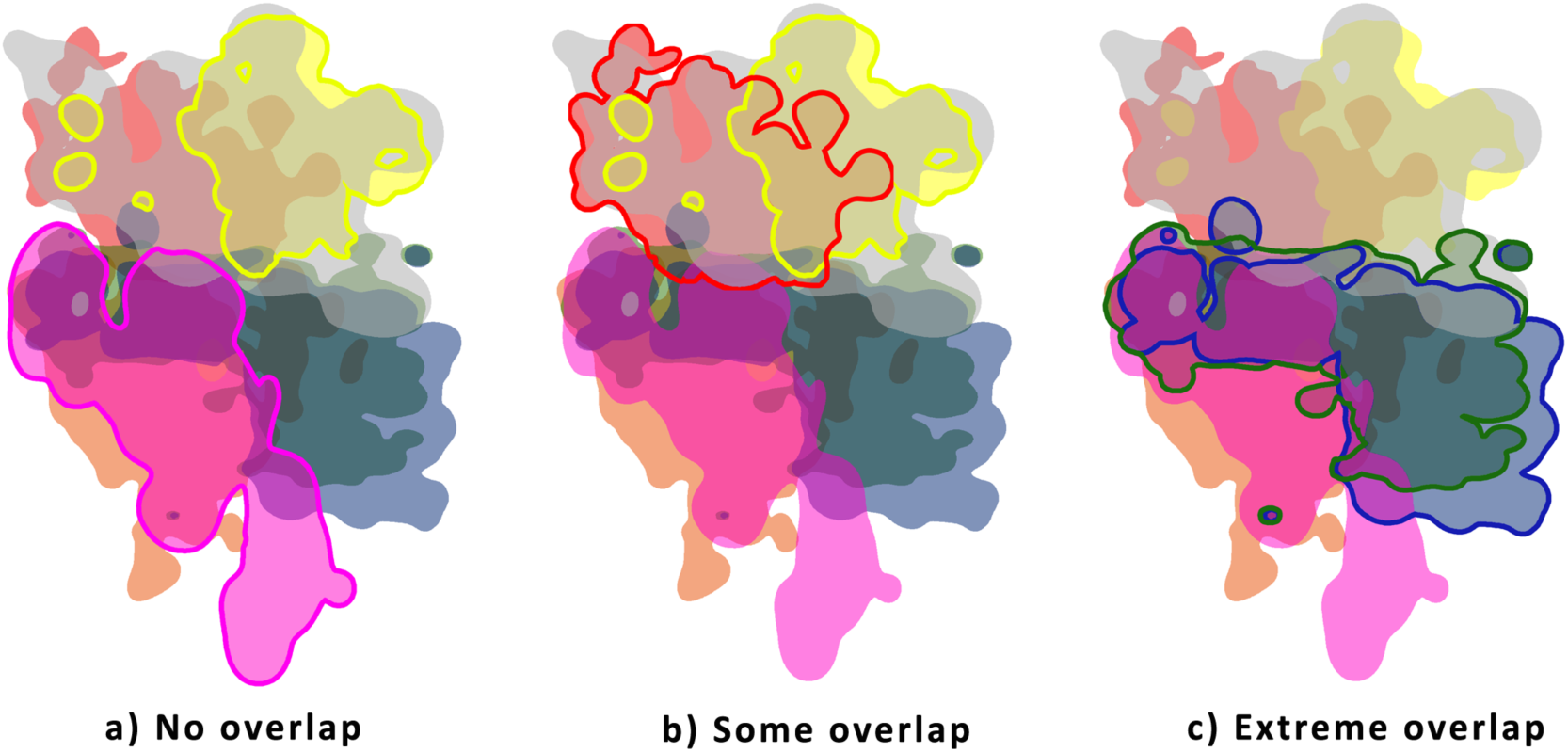
Examples of home range overlap among pairs of individuals within the Mangevo ruffed lemur community: a) Females Pink-Yellow and Radio-Yellow have no spatial overlap among home ranges (i.e., UDOI = 0.00), whereas b) females Radio-Yellow and Radio-Red and c) females Radio-Blue and Radio-Green share some (UDOI = 0.106) to nearly complete home range overlap (UDOI = 0.951), respectively.

**Table 1.** Summary of association indices, utilization distribution overlap indices (UDOIs), and relatedness (*r*) overall and by reproductive season.

| Association Indices | n | Mean | SD | Min | Max |
| --- | --- | --- | --- | --- | --- |
| Overall | 136 | 0.05 | 0.03 | 0.00 | 0.80 |
| Nonreproductive | 136 | 0.04 | 0.12 | 0.00 | 0.82 |
| Mating/Gestation | 105 | 0.06 | 0.13 | 0.00 | 0.79 |
| Birth/Lactation | 91 | 0.07 | 0.15 | 0.00 | 0.82 |
| UDOIs | n | Mean | SD | Min | Max |

| Overall | 136 | 0.21 | 0.36 | 0.00 | 1.90 |
| --- | --- | --- | --- | --- | --- |
| Nonreproductive | 136 | 0.17 | 0.27 | 0.00 | 1.97 |
| Mating/Gestation | 120 | 0.15 | 0.29 | 0.00 | 1.74 |
| Birth/Lactation | 78 | 0.20 | 0.43 | 0.00 | 2.13 |
| Relatedness | n | Mean | SD | Min | Max |
| Overall | 171 | -0.06 | 0.02 | -0.72 | 0.65 |
| Female-female | 45 | -0.13 | 0.04 | -0.60 | 0.52 |
| Female-male | 90 | -0.15 | 0.02 | -0.72 | 0.28 |
| Male-male | 36 | -0.13 | 0.04 | -0.54 | 0.60 |

### Relatedness

Genotypes were 93% complete; all subjects (n = 38) were scored for at least 12 loci (average = 14, range = 12 to 15). Allelic richness was 4.33 and average observed heterozygosity was 0.400. There were no significant deviations from Hardy-Weinberg Equilibrium for any of the loci examined, nor was there evidence of null alleles.

Results of the rarefaction curve (y = 0.7991, r^2^ =0.9992) showed average relatedness values stabilizing after 5 loci, with the difference between mean relatedness using 5 and 6 loci changing by only 0.95% (0.023), and the difference between using 6 and 7 loci changing by only 0.56% (0.016). Thus, subsequent dyadic r-value calculations included all possible dyads (n = 703 dyads), as all individuals could be compared at 5 or more loci. Average pairwise relatedness among adults within the community was −0.06 ± 0.02 and ranged from −0.72 to 0.65 (Table 1; Figure 2).

**Figure 2.**
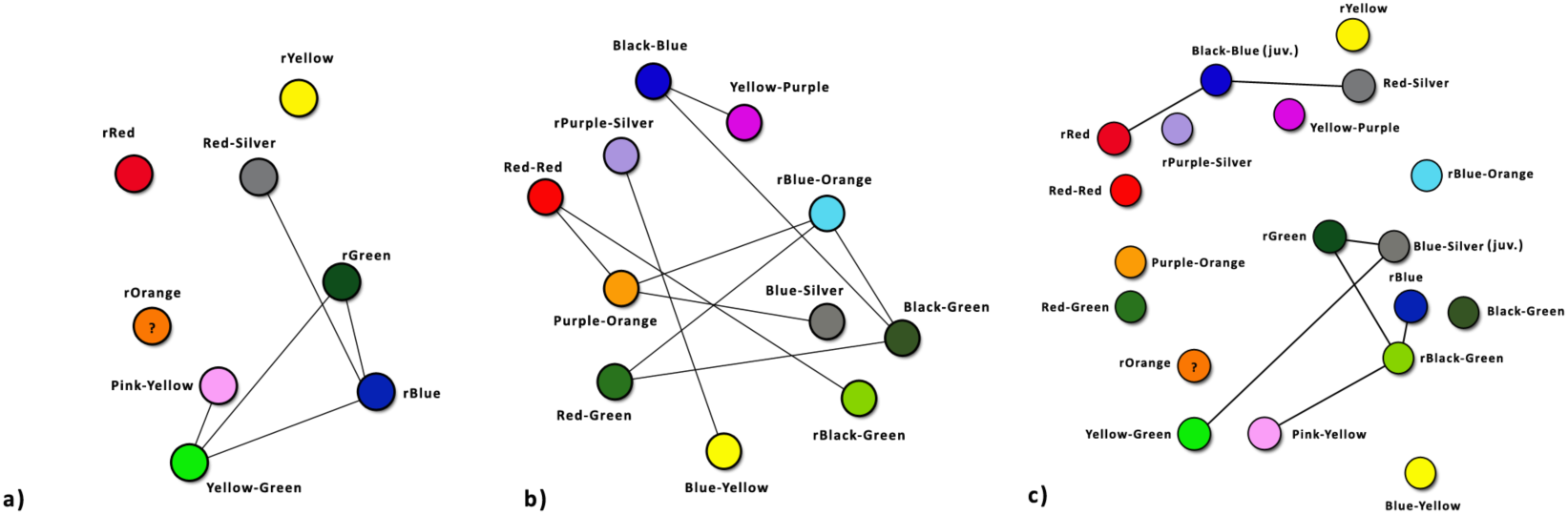
Pairwise genetic relatedness (r ≥ 0.25) among adult a) female-female dyads, b) male-male dyads, and c) female-male dyads within the Mangevo social community. “?” indicates individuals for which genotype data are unavailable. Nodes are organized according to individual home range centroids.

Fewer than ten percent (8.8%) of dyads within the community were genetic relatives. Despite this, more than three-quarters of adult females (n=6 of 7 for which genetic data were available; 85.7%) and all of adult males (n=11 of 11; 100%) were related (r ≥ 0.25) to at least one, and up to as many as three other same-sex relative(s) within the community (Figure 2).

### Relationships among social associations, spatial overlap, and relatedness

Using the full dataset, Mantel tests revealed that kinship was unrelated to either home range overlap (UDOI; n = 25, r = 0.047 p = 0.724) or association indices (AI; n = 14, r = 0.120, p = 0.294), whereas UDOI and AI were significantly correlated (n = 17, r = 0.789, p << 0.001; Figure 3). This pattern held across all three reproductive seasons (Table 2). It is worth noting, however, that the correlation coefficient between UDOI and AI in the nonbreeding season was much lower, albeit still significantly positive, than the mating/gestation and birth/lactation seasons, which were very similar (Table 2). Similarly, when we used MRQAPs to jointly analyze the effects of UDOI, kinship, and sex on AI, in every time period we analyzed, the partial correlations between UDOI and AI were always significantly positive, while kinship and sex were never significantly correlated with AI (Table 3). These results support prediction P1.1, but do not support prediction P1.2.

**Figure 3.**
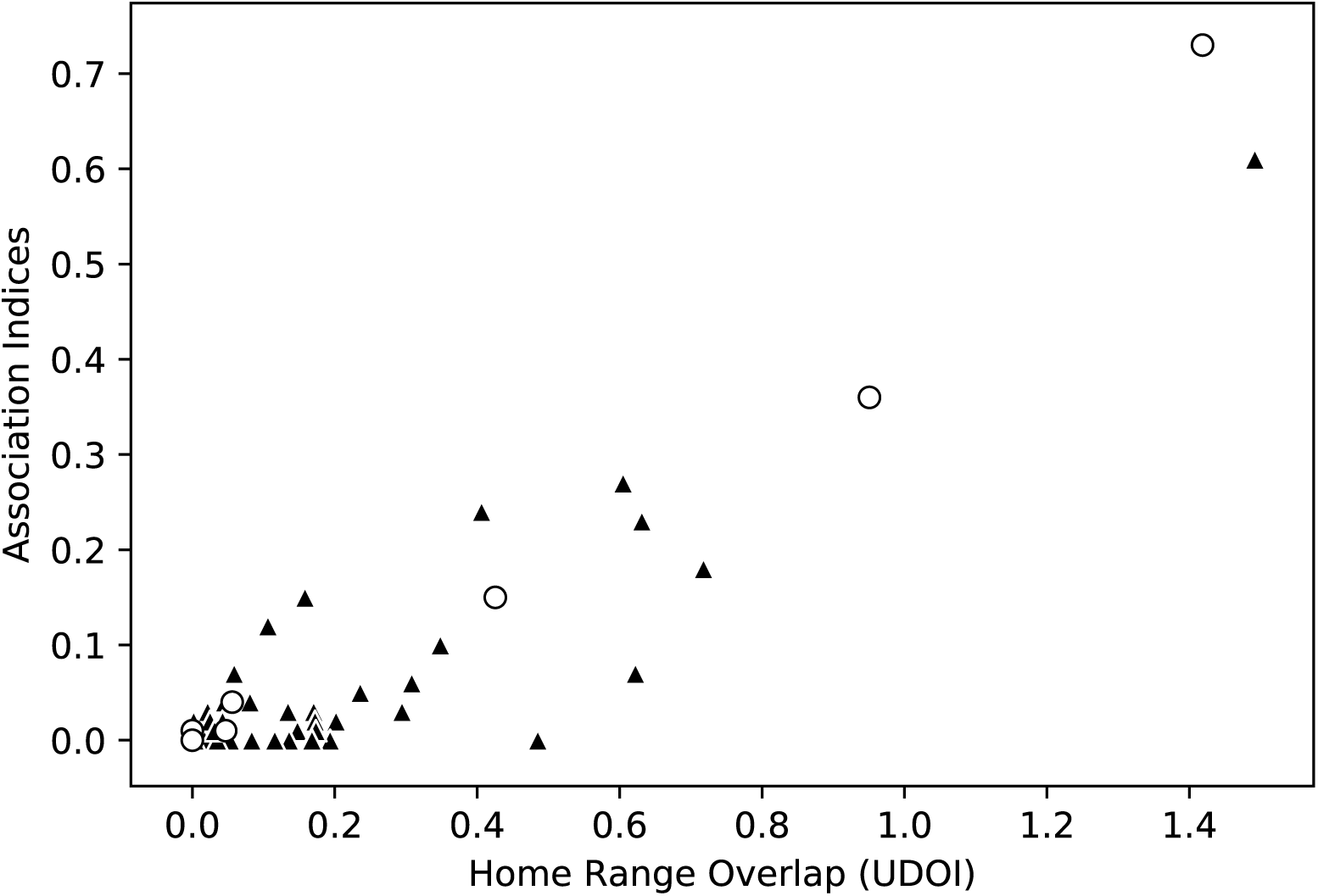
Scatterplot illustrating the relationship between social association (AI) and home range overlap (UDOI). AIs range from 0 (no association) to 1 (100% association). Similarly, UDOIs range from 0 (no overlap) to 1 (complete overlap); values of >1 are possible if both UDs are nonuniformly distributed and also have a high degree of overlap. Values <1 indicate less overlap relative to uniform space, whereas values >1 indicate higher than normal overlap relative to uniform space. White circles indicate kin (r ≥ 0.25); black triangles indicate non-kin (r<0.25).

**Table 2.** Results from Mantel tests correlating Association Indices with utilization distribution overlap indices (UDOIs), and relatedness overall and by reproductive season.

| Season | Variable | Records | N | Correlation | p-value |
| --- | --- | --- | --- | --- | --- |
| Overall | UDOI | 11,171 | 17 | 0.789 | <<0.001 |
| Overall | Relatedness | 11,171 | 14 | 0.120 | 0.294 |
| Mating | UDOI | 3,267 | 13 | 0.891 | <<0.001 |
| Mating | Relatedness | 3,267 | 13 | 0.099 | 0.392 |
| Nonbreeding | UDOI | 5,051 | 14 | 0.566 | <<0.001 |
| Nonbreeding | Relatedness | 5,051 | 14 | 0.149 | 0.184 |
| Birth/gestation | UDOI | 2,664 | 11 | 0.892 | <<0.001 |
| Birth/gestation | Relatedness | 2,664 | 11 | 0.159 | 0.254 |

Note: Overall includes 2007-2008, whereas seasonal measures are from 2008 only. N indicates the number of individuals included in each analysis; individuals had to be represented in both datasets in order to be included in the analyses for a given season.

**Table 3.** Results from a multiple regression quadratic assignment procedure (MRQAP) investigating the partial relationships of association (AI) with utilization distribution overlap indices (UDOI), relatedness, and sex.

| Season | N | UDOI |  | Relatedness |  | Sex |  |
| --- | --- | --- | --- | --- | --- | --- | --- |
|  |  | Partial<br>corr | *p*-value | Partial<br>corr | *p*-value | Partial<br>corr | *p*-value |
| Overall | 14 | 0.920 | <<0.001 | 0.035 | 0.754 | 0.056 | 0.617 |
| Nonbreeding | 11 | 0.655 | <<0.001 | 0.123 | 0.356 | -0.113 | 0.460 |
| Mating/gestation | 10 | 0.873 | <<0.001 | -0.049 | 0.742 | 0.281 | 0.720 |
| Birth/lactation | 8 | 0.941 | <<0.001 | 0.296 | 0.144 | 0.102 | 0.530 |

### Relatedness among preferred associates

We identified 18 (of 136 total) pairs of preferred associates, i.e., dyads with AIs that were significantly higher than other dyads in the community. Of these preferred associates, we were able to determine relatedness for 11 (91 total dyads with data for association and relatedness). Our logistic regression model revealed that relatedness did not predict whether a pair of individuals became preferred associates (coefficient = 0.087, p = 0.41) and thus did not support prediction P2.1.

## Discussion

Results from this study illustrate complex relationships among social association, space use, and kinship in wild black-and-white ruffed lemurs, patterns that—while unusual for primates—are well aligned with much of the broader mammalian literature. We found that ruffed lemur social associations varied immensely, ranging from no association between some individuals to dyads observed together more than 75% of the time. From a community perspective, the social network was sparse, with weak associations (AIs) being common. Similarly, home range overlap was minimal and average relatedness among community members was low. These patterns were consistent throughout the year and across reproductive seasons. Together, these and earlier lines of evidence (e.g., Baden, 2011; Baden et al., 2016) can be used to characterize ruffed lemurs as having a spatiotemporally dispersed fission-fusion social organization with weak social and kinship ties.

Kin selection theory (Hamilton, 1964) has long been invoked to explain the social preferences observed among mammals (e.g., Archie et al., 2006; Frère et al., 2010; Godde et al., 2015; Smith, 2014; Wahaj et al., 2004), particularly in primates (reviewed in Langergraber, 2012; Silk, 2002). In this study, however, we found no evidence that kinship structured either spatial overlap or social association overall in the community. In contrast to other fission-fusion species exhibiting similar patterns (Carter et al. 2013; Best et al. 2014), we further found that relatedness did not predict social preference. Instead, the closest social ties in the community appear to occur primarily between females and their preferred—and often unrelated—male social partners, followed by mothers and their pre-dispersal-aged subadult and adult offspring. Nevertheless, although fewer than ten percent (8.8%) of adult dyads within the community were genetic relatives, nearly three-quarters of adult females (85.7%) and all of adult males (100%) were closely related to at least one, and up to as many as three other same-sex relative(s). Thus, while our relatedness estimates suggest that ruffed lemur communities include both kin and non-kin, and that preferred associates are sometimes close relatives, kin are not forming spatial or social networks across the larger communal range. These patterns contrast with the spatially structured matrilines described in *Microcebus murinus* (Eberle & Kappeler, 2006; Radespiel, Lutermann, Schmelting, Bruford, & Zimmermann, 2003), the only other communally breeding strepsirrhine. Some authors have used this model to hypothesize that spatially structured kin networks have, at least in part, facilitated the evolution of cooperative infant care (Eberle & Kappeler, 2006). Cooperative or communal breeding is exceedingly rare among primates (Tecot et al. 2013). Only three nonhuman primates engage in this behavior: callitrichines (marmosets and tamarins), cheirogaleids (mouse lemurs), and *Varecia* (Baden, Raboin, Tecot, n.d.). Like cheirogaleids, callitrichines also rely on close relatives to share the burden of infant care (reviewed in Erb & Porter 2017). *Varecia* are the only group that does not rely on close relatives for care (Baden et al., 2013), thereby distinguishing *Varecia*’s infant care strategies from other cooperatively breeding nonhuman primates.

These results build on growing evidence that space use is an important predictor of social association—perhaps moreso than kinship—particularly in taxa characterized by high fission-fusion dynamics (Best et al., 2014; K. D. Carter et al., 2013; Frère et al., 2010; Strickland et al., 2014). Indeed, for two individuals to interact they must be in relatively close physical proximity (Farine 2015). For most animals, this means that social associations are therefore shaped by their movement decisions (Emlen and Oring 1977; Farine et al. 2016; Bonnell et al. 2017), which are, in turn, influenced by aspects of their physical environment (e.g., habitat structure, natural and/or anthropogenic barriers, resource availability and distribution; reviewed in He et al. 2019). Among these environmental variables, resource availability and distribution play a fundamental role in shaping the space use of animals (e.g., Brown and Orians 1970; Clutton-Brock and Harvey 1977) and by extension, their patterns of inter-individual spatial proximity (e.g., Minta 1992; Croft et al. 2011) and social relationships (e.g., Sterck et al. 1997; Bejder, Fletcher & Brager 1998; Wolf et al. 2007; Carter et al. 2009).

To this point, we found that while home range overlap explained most of the variation seen in social association in ruffed lemurs, some variation remained unaccounted for, suggesting that other social, ecological, and biological factors must also be at play. However, it is possible that even weak or infrequent social associations may facilitate cooperative resource defense against other frugivorous competitors. For instance, ruffed lemurs actively defend fruit-bearing trees against larger brown lemur (Genus *Eulemur*) groups for days or even weeks during the resource scarce austral winter (Baden, personal observation). Perhaps social association during these times better equips otherwise solitary individuals to defend valuable fruit resources against interspecific competitors. Under this scenario, we would predict higher AIs during resource scarce seasons, periods that correspond primarily with mating/gestation, but also birth/lactation seasons. However, AIs were lowest in the nonbreeding, resource abundant periods (though not significantly so), lending minimal support for this hypothesis. It is possible, however, that home range overlap as measured by UDOIs was insufficient to characterize the spatiotemporal component of associations at highly contested resources. Future analyses may therefore benefit from a more nuanced investigation of the spatially explicit role valuable resources play in the social associations of this species.

In addition, communal breeding plays an important role in female reproductive success in this species (Baden et al., 2013) and might also be important in driving social preferences. For instance, recent work suggests social networks in guppies may be structured by the propensity for non-kin to cooperate (Croft et al., 2009), which could lead to the maintenance of cooperation in the absence of kin assortment (Fletcher & Doebeli, 2009). These lines of research offer exciting opportunities to better understand the myriad factors shaping social preferences in fission-fusion species.

Nevertheless, shared space use does not always necessitate social association. In this study, we found that not all dyads with a high degree of home range overlap were close social associates. Indeed, many dyads with nonzero spatial overlap were never observed together. Similar patterns have been observed in dolphins, giraffes, and water dragons, wherein subjects did not associate, despite sharing complete or near complete home range overlap (i.e., ‘social avoidance’ in dolphins: Frère et al., 2010; giraffes: K. D. Carter et al., 2013; water dragons: Strickland et al., 2014, 2017). In such cases, individuals may actively and consistently avoid one another, despite the spatio-temporal potential for repeated interactions (reviewed in Strickland et al. 2017). Like predator avoidance, social avoidance can be costly. Thus, in future studies, it will be critical to simultaneously consider the consequences of both social attraction and avoidance when studying the evolution of sociality.

Together, these and earlier results raise important questions related to causation. What motivates social association? Are individuals that bias their time toward overlapping areas simply more likely to associate? Or, in cases where patterns of spatial overlap and social association do not align, is there some additional force shaping these spatial and social decisions? Evaluating these and other alternatives require further investigation.

## Acknowledgements

Research would not have been possible without the logistical support of The Institute for the Conservation of Tropical Environments (ICTE), MICET and Centre ValBio, or permission from ANGAP/Madagascar’s National Parks. Drs. Randy Junge, Felicia Knightly, Angie Simai, Edward E. Louis, Jr., and staff of the Madagascar Biodiversity Project and Prosimian Biomedical Survey Project provided invaluable assistance during capture procedures. Solo Justin, Telo Albert, Lahitsara Pierre, Razafindrakoto Georges, Leroa, Velomaro, Reychell Chadwick, Lindsay Dytham, and A.J. Lowin assisted with data collection and provided invaluable support and advice on the ground, while Gary Aronsen provided support in the Yale Molecular Anthropology Lab. We thank two anonymous reviewers, as well as James Higham, for their insightful comments on earlier versions of this manuscript. We are further grateful for resources and support from the Center for High Performance Computing at the University of Utah. Funding was generously provided by National Science Foundation DDIG (BSC-0725975), the US Department of State Fulbright Fellowship, The Leakey Foundation, Primate Conservation, Inc., Conservation International’s Primate Action Fund, Stony Brook University, Henry Doorly Zoo, and Yale University.

## Notes

#### Summary of Updates

New analyses; text revisions; title modified

